# Partitioning Genotypic and Environmental Variance of Fresh Root Yield in Provitamin A Cassava Genotypes Using Mixed Models

**DOI:** 10.1101/2025.11.18.688222

**Authors:** Olusegun Badewa, Elizabeth Parkes, Andrew Gana, Eli Tsado, Kehinde Tolorunse, Peter Iluebbey, Patrick Akpotuzor, Toye Ayankanmi

## Abstract

This study employed a multi-model statistical approach to evaluate the yield performance and environmental responsiveness of 42 provitamin A cassava genotypes across two contrasting cropping seasons. Four linear mixed models spanning BLUP-based random effects, fixed-effect genotype comparisons, genotype-by-environment interaction analysis, and variance partitioning were used to dissect phenotypic variation in fresh root yield. Months After Planting (MAP) emerged as the most influential factor, accounting for 19.82% of total variance, while genotypic effects were modest (H² = 4.4%) and seasonal effects reached statistical significance when accessions were treated as fixed. Residual variance dominated (70.98%), suggesting the influence of unmeasured factors such as soil properties, pest pressure, and microclimatic variation. Rainfall data revealed a stark contrast between seasons-127.38 mm in 2019/2020 versus 8.45 mm in 2020/2021 highlighting an environmental stress pattern that shaped genotype performance. Accessions IITA-TMS-IBA980581(WChk-White Check), IITA-TMS-IBA180017, and IITA-TMS-IBA180146 maintained high yields across both seasons, indicating broad adaptability, while IITA-TMS-IBA180182 and IITA-TMS-IBA180065 performed better under low-moisture conditions, suggesting drought responsiveness. These findings underscore the importance of integrating rainfall patterns and genotype-specific environmental sensitivity into cassava breeding pipelines. Future multilevel models incorporating locations and climatic data could enhance selection precision and support the development of resilient varieties for variable agroecologies.

## Introduction

Cassava (*Manihot esculenta* Crantz) is a staple root crop widely cultivated across sub-Saharan Africa for its resilience to marginal soils and its role in food security (Sakadzo et al., 2025; Scaria et al., 2023). In recent years, breeding efforts have intensified to develop biofortified cassava genotypes rich in provitamin A, addressing both nutritional deficiencies and agronomic performance (Delgado et al., 2024). However, the productivity of cassava genotypes is highly influenced by environmental factors, including seasonal rainfall patterns, soil conditions, and harvest timing (Phanthanong et al., 2025).

Understanding genotype × environment (G×E) interactions is critical for identifying stable and high-yielding genotypes across diverse agroecological zones (Begna, 2020). Yield variability in cassava is often shaped by both genetic potential and environmental responsiveness, particularly under fluctuating climatic conditions (Phanthanong et al., 2025). Months After Planting (MAP) is a key developmental metric that influences root bulking, moisture availability, and overall yield outcomes (Badewa et al., 2020). Evaluating genotypic performance across multiple MAP stages and seasons provides insights into temporal stability and environmental adaptation (Okoma et al., 2025).

This study aimed to assess the fresh root yield (FYLD) of 42 provitamin A cassava genotypes across two contrasting cropping seasons (2019/2020 and 2020/2021) and three harvest periods (6, 9, and 12 MAP). Using a split-plot design and mixed-effects modeling, we quantified variance components, ranked genotypes using BLUP estimates, and explored G×E interactions to identify stable performers under variable rainfall conditions. The findings contribute to cassava breeding strategies by highlighting genotypes with consistent yield potential and resilience to seasonal stress.

## Materials and Methods

### Study Location and Plant Materials

The field evaluation was conducted over two consecutive cropping seasons (2019/2020 and 2020/2021) at the International Institute of Tropical Agriculture (IITA), Ibadan, Nigeria (Latitude 7.488249°N, Longitude 3.904875°E; altitude: 207 m). The site is representative of the humid tropics and has historically served as a core location for cassava breeding trials. A total of 42 cassava genotypes rich in provitamin A were selected for evaluation. These included three white-fleshed checks (TMEB419, TME693, and IBA980581) and one yellow-fleshed check (IITA-TMS-IBA070593), all sourced from the Cassava Breeding Unit at IITA.

### Experimental Design and Data Collection

This study evaluated the yield performance of 42 provitamin A cassava genotypes across two cropping seasons (2019/2020 and 2020/2021) and three harvesting stages, measured as Months After Planting (MAP): 6MAP (December), 9MAP (March), and 12MAP (June). The experiment was structured as a split-plot design with two replications, where genotype served as the main plot factor and harvest time (MAP) as the subplot factor. Blocks were represented by the factor REP, and trials were conducted under varying seasonal conditions to capture environmental responsiveness.

Within each replication, genotypes were randomly distributed across the three harvest times to minimize positional bias. Each plot measured 4 m × 2 m (8 m²) and was subdivided into three equal subplots of 2.67 m², each containing four plants arranged in two rows. Plant spacing was maintained at 1 m × 0.8 m, resulting in a total experimental area of 336 m² per replicate and 672 m² across both replications.

Only fresh root yield (FYLD) was evaluated. At each harvest time, all plants within each subplot were harvested, and total fresh root weight was recorded using a calibrated digital scale. Yield data were expressed in kilograms per plot and later converted to tons per hectare (t/ha) for analysis.

### Agronomic Management

Standard agronomic practices were followed throughout the trial. Manual weeding was conducted as needed, and no supplemental irrigation or fertilizer was applied to simulate typical farmer-managed conditions. Pest and disease monitoring was performed regularly, though no major outbreaks were recorded during the study period.

### Statistical Analyses

To dissect the effects of genotype, seasonality, and developmental timing on yield, four complementary linear mixed models were fitted using R (version 4.3.2) (R Core Team, 2023). The models incorporated the factors MAP, Seasons, REP, and Accession_name. Fixed effects were evaluated using Type III ANOVA with Satterthwaite’s approximation for degrees of freedom.

Model 1: A mixed-effects model (lmer) treating *Accession_name* as a random effect, with *MAP*, *Seasons*, and *REP* as fixed effects, including their interactions. This model estimated genotypic variance and BLUPs while testing for MAP × Season effects (Bates et al., 2015).

Model 2: A fixed-effects model (lm) specifying *Accession_name* as a categorical fixed factor to assess individual genotype effects in relation to *MAP*, *Seasons*, and *REP*.

Model 3: A genotype-by-environment interaction model (lmer) including *MAP × Seasons × Accession_name* as fixed effects, used to detect crossover interactions and season-specific genotype responsiveness.

Model 4: A variance partitioning model (lmer) with *MAP*, *Seasons*, and *Accession_name* and REP treated as random effects, to quantify the relative contributions of genetic, seasonal, and management factors to total phenotypic variance.

Variance components were extracted using VarCorr, and heritability (H²) was computed as the ratio of genotypic variance to total phenotypic variance. Diagnostic checks were performed using the performance package, and estimated marginal means were computed using emmeans (Lenth, 2022).

## RESULTS

### Model 1

**Table 1.1:** ANOVA Summary for Mixed-effects model evaluating fresh root yield across environments.

| Effect | F value | Pr(>F) | Significance | Interpretation |
| --- | --- | --- | --- | --- |
| MAP | 92.540 | p< 0.001 | *** | Strong influence of MAP on yield |
| Seasons | 2.660 | 0.103 | ns | Season alone doesn't significantly affect FYLD |
| REP | 0.000 | 0.983 | ns | No significant effect from replication |
| MAP × Seasons | 0.179 | 0.672 | ns | No significant interaction effect |

\*\*\* $p < 0.001$ , \*\* $p < 0.01$ , \* $p < 0.05$ , $p < 0.1$ , ns = not significant

#### Harvest Timing (MAP) as the Primary Driver of Yield Variation

The analysis revealed a highly significant effect of harvest timing (MAP), indicating that yield was strongly influenced by the number of months after planting. In contrast, seasonal variation and replication effects were not statistically significant, suggesting that yield performance remained relatively consistent across seasons and replicates. Additionally, the interaction between MAP and Season was non-significant, implying that the influence of harvest timing on yield did not vary meaningfully between seasons. These findings underscore the dominant role of MAP in shaping cassava productivity, while other environmental factors exerted minimal influence in this model (Table 1.1).

#### Genotypic Yield Rankings Based on BLUP Estimates

As presented in Table 1.2, the BLUP-estimated fresh root yield values for cassava genotypes, highlighting the top five and bottom five performers across all environments. Genotype IITA-TMS-IBA180146 exhibited the highest predicted yield (BLUP: 4.97), followed by IITA-TMS-IBA180081 and the white-fleshed check IITA-TMS-IBA980581(WChk), indicating strong and consistent productivity. In contrast, genotypes such as IITA-TMS-IBA180031 (BLUP: -3.92) and IITA-TMS-IBA180173 (BLUP: -3.35) showed the lowest predicted yields, suggesting poor adaptation or performance under the tested conditions. These BLUP estimates provide a foundational ranking of genotypic potential, serving as a benchmark for subsequent stability and interaction analyses.

**Table 1.2:** BLUPs for Genotypic Performance for Fresh Root Yield.

| Top 5 High Performers | Bottom 5 Performers: |
| --- | --- |
| IITA-TMS-IBA180146 (BLUP: 4.97) | IITA-TMS-IBA180031 (BLUP: -3.92) |
| IITA-TMS-IBA180081 (BLUP: 3.50) | IITA-TMS-IBA180173 (BLUP: -3.35) |
| IITA-TMS-IBA980581 (WChk) (BLUP: 3.22) | IITA-TMS-IBA180158 (BLUP: -3.32) |
| IITA-TMS-IBA180210 (BLUP: 2.65) | IITA-TMS-IBA180294 (BLUP: -2.44) |
| IITA-TMS-IBA180073 (BLUP: 2.24) | IITA-TMS-IBA180271 (BLUP: -2.45) |

#### Environmental Influence Overshadows Genetic Contribution to Yield

From the variance components derived from the mixed-effects model for fresh root yield as shown in Table 1.3. The genotypic variance (σ²g = 10.54) indicates modest genetic differentiation among accessions, while the residual variance (229.30) accounts for the majority of yield variation, suggesting strong environmental influence or unexplained variability. The total phenotypic variance (σ²p = 239.85) reflects the overall variation observed in the dataset. The estimated broad-sense heritability (H² = 0.044) reveals that only 4.4% of the yield variation is attributable to genetic factors, underscoring the dominant role of environmental conditions in shaping cassava productivity under the tested conditions.

**Table 1.3:** Variance Component of Model 1.

| Component | Value | Interpretation |
| --- | --- | --- |
| Genotypic Variance($\sigma^2g$) | 10.54 | There's some genetic differentiation between accessions, but it's small. |
| Residual Variance | 229.30 | Most of the variation in yield is due to environmental factors or unexplained noise. |
| Phenotypic Variance($\sigma^2p$) | 239.85 | Total variation observed in FYLD. |
| Heritability ($H^2$) | **0.044** | Only **4.4%** of the yield variation is attributable to genetic factors. |

### Model 2

**Table 2.1:**
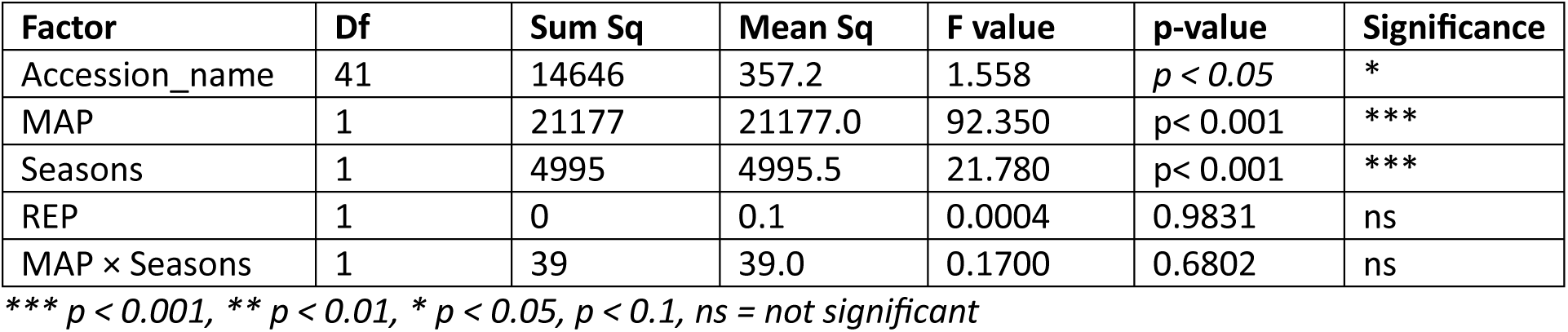
ANOVA Table for Fixed Effect Model.

| Factor | Df | Sum Sq | Mean Sq | F value | p-value | Significance |
| --- | --- | --- | --- | --- | --- | --- |
| Accession_name | 41 | 14646 | 357.2 | 1.558 | $p < 0.05$ | * |
| MAP | 1 | 21177 | 21177.0 | 92.350 | $p < 0.001$ | *** |
| Seasons | 1 | 4995 | 4995.5 | 21.780 | $p < 0.001$ | *** |
| REP | 1 | 0 | 0.1 | 0.0004 | 0.9831 | ns |
| MAP × Seasons | 1 | 39 | 39.0 | 0.1700 | 0.6802 | ns |

\*\*\* $p < 0.001$ , \*\* $p < 0.01$ , \* $p < 0.05$ , $p < 0.1$ , ns = not significant

#### Yield Variation Driven by MAP and Genotype with Seasonal Influence

Table 2.1 presents the ANOVA results from the fixed-effects model used to assess individual genotype contributions to fresh root yield. The factor Accession_name was statistically significant (F = 1.558, p = 0.0176), indicating that genotypic differences contributed meaningfully to yield variation. As in Model 1, MAP (Months After Planting) remained highly significant (F = 92.35, p < 0.001), confirming its dominant influence on productivity. Season also showed a strong effect (F = 21.78, p <0.05), suggesting that seasonal conditions impacted genotype performance more clearly in this model. Replication and the MAP × Season interaction was not significant, indicating consistency across replicates and no differential seasonal response to harvest timing. Overall, this model highlights the importance of genotype, harvest timing, and seasonal conditions in shaping cassava yield outcomes.

#### Cross-Model Validation of Genotypic Performance Patterns

Table 2.2 compares fixed-effect estimates from Model 2 with BLUP values from Model 1 for selected cassava genotypes. Genotype IITA-TMS-IBA180146 consistently ranked as a top performer across both models, with a high fixed estimate (10.78) and BLUP value (4.97), though its p-value (0.0819) was marginal. In contrast, genotypes IITA-TMS-IBA180031, IITA-TMS-IBA180158, and IITA-TMS-IBA180173 were identified as significant underperformers, each showing strongly negative fixed estimates and BLUP values, with p-values below 0.05. This comparison highlights the alignment between fixed and mixed model outputs, reinforcing the reliability of BLUP-based selection while confirming statistical significance for low-yielding genotypes (Figure 2.1; 2.2)

**Table 2.2:** Accession Fixed Estimates vs. BLUPs.

| Accession Name | Fixed Estimate | BLUP Value | p-value | Performance Summary |
| --- | --- | --- | --- | --- |
| IITA-TMS IBA180146 | 10.78 | 4.97 | 0.0819 | Top performer across both models |
| IITA-TMS-IBA180031 | -14.23 | -3.92 | 0.0218 | Significant underperformer |
| IITA-TMS-IBA180158 | -12.53 | -3.32 | 0.0433 | Significant underperformer |
| IITA-TMS-IBA180173 | -12.62 | -3.35 | 0.0417 | Significant underperformer |

**Figure2.1:**
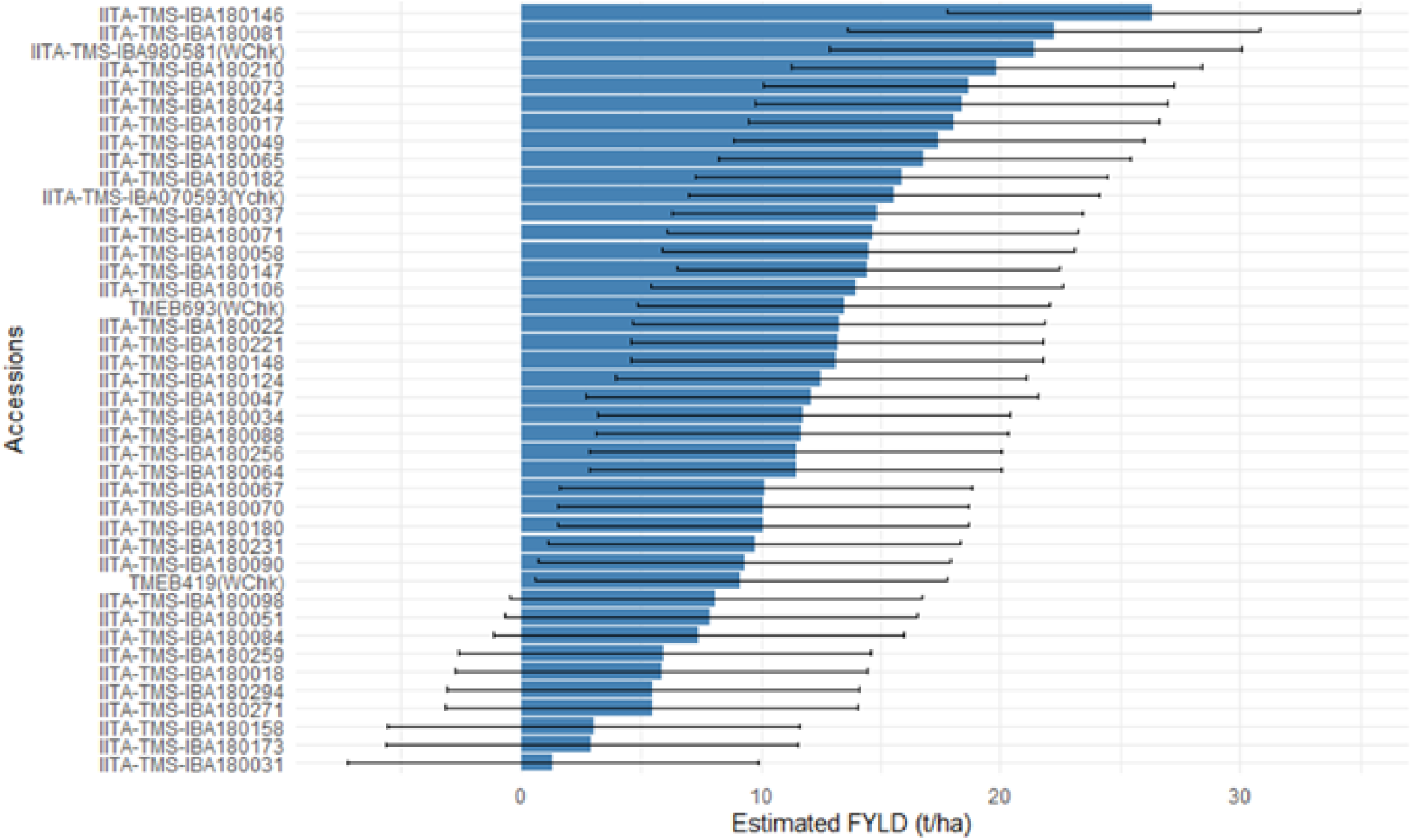
Estimated Genotype Effects (Fixed Model)

**Figure 2.2:**
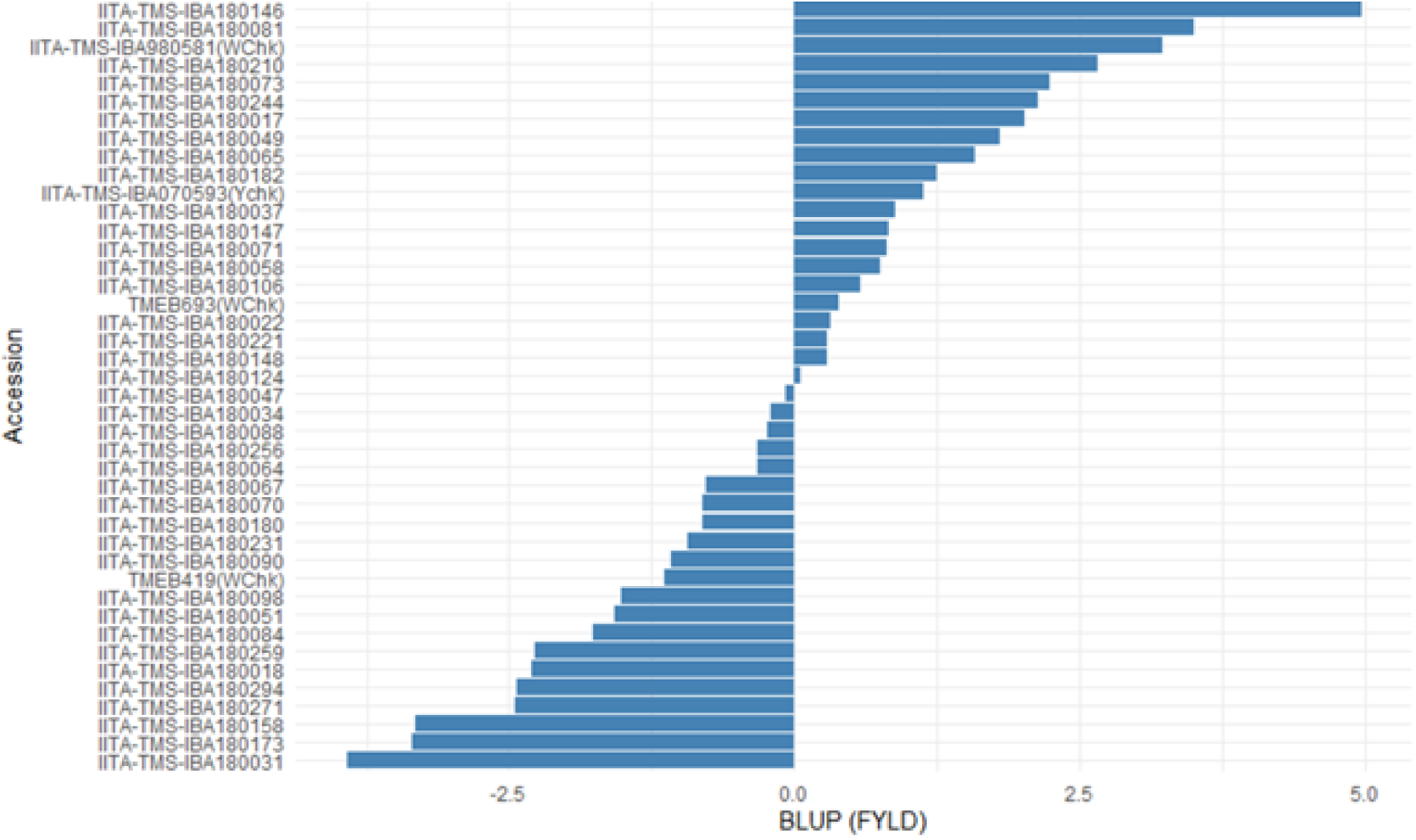
Genotype Ranking by BLUPs.

### Model 3

**Table 3.1:**
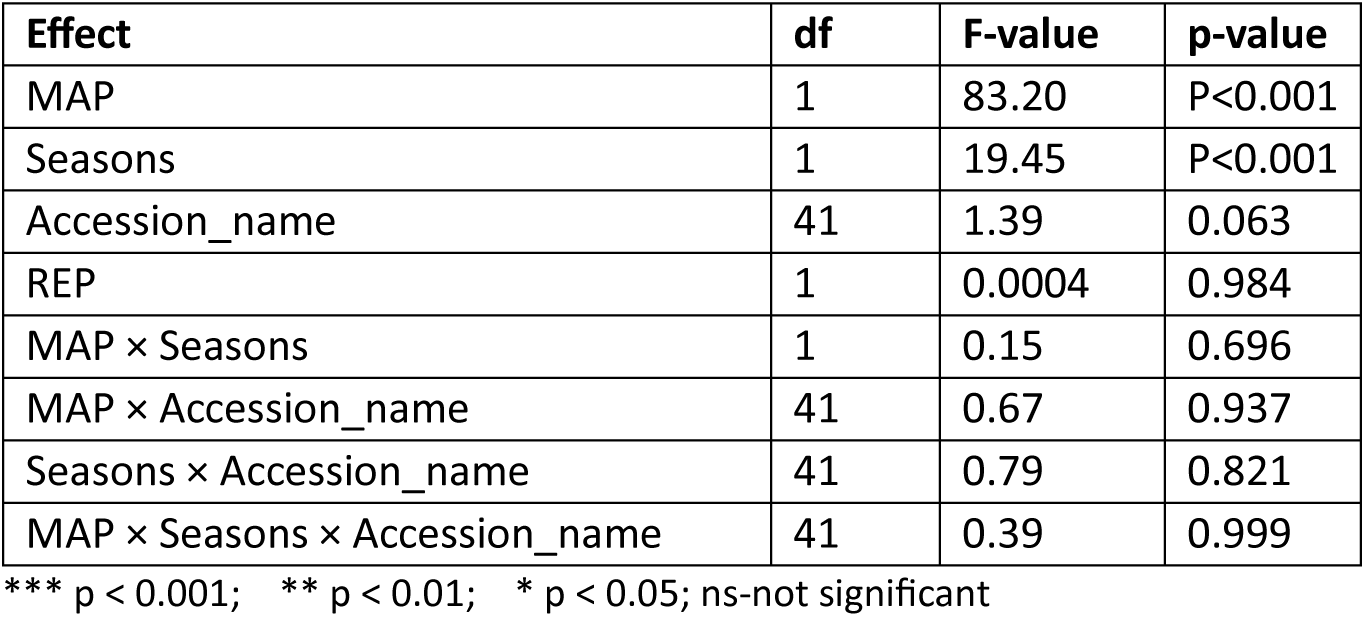
ANOVA Table Showing Main and Interaction Effects of MAP, Seasons, and Genotypes on Yield Performance.

| Effect | df | F-value | p-value |
| --- | --- | --- | --- |
| MAP | 1 | 83.20 | P<0.001 |
| Seasons | 1 | 19.45 | P<0.001 |
| Accession_name | 41 | 1.39 | 0.063 |
| REP | 1 | 0.0004 | 0.984 |
| MAP × Seasons | 1 | 0.15 | 0.696 |
| MAP × Accession_name | 41 | 0.67 | 0.937 |
| Seasons × Accession_name | 41 | 0.79 | 0.821 |
| MAP × Seasons × Accession_name | 41 | 0.39 | 0.999 |

\*\*\* p < 0.001; \*\* p < 0.01; \* p < 0.05; ns-not significant

#### Effect of Genotype, Season, and MAP on Cassava Yield Performance

The analysis of variance revealed that MAP (Months After Planting) and Seasons had statistically significant effects on yield performance, with F-values of 83.20 and 19.45 respectively, and p-values less than 0.001. These results indicate that variation across MAPs and seasons strongly influenced genotype performance.

The effect of Accession_name (genotype) was marginally non-significant (F = 1.39, p = 0.063), suggesting limited variation among genotypes in their overall mean yield across environments. Replication (REP) had no significant effect (F = 0.0004, p = 0.984), confirming consistency across experimental blocks.

None of the interaction terms-including MAP × Seasons, MAP × Accession_name, Seasons × Accession_name, and the three-way interaction-were statistically significant (all p > 0.69), indicating that genotype performance did not vary significantly across combinations of location and season (Table 3.1).

#### Seclection based on BLUPS and Standard Deviation of BLUPS

The analysis evaluated cassava genotypes using Best Linear Unbiased Predictions (BLUPs) to estimate yield potential, alongside the standard deviation of BLUPs (BLUP_SD) to assess yield stability across seasons. Genotypes were ranked based on their BLUP values, with higher scores indicating superior predicted performance. BLUP_SD values ranged from 0.33 to 1.67, reflecting varying degrees of environmental sensitivity. Genotypes such as *IITA-TMS-IBA180146* and *IITA-TMS-IBA180210* exhibited high BLUPs (4.97 and 2.65, respectively) with low BLUP_SD values (0.42 and 0.39), indicating both strong yield potential and consistent performance. In contrast, genotypes like *IITA-TMS-IBA180182* and *IITA-TMS-IBA180073* showed relatively low BLUPs (1.25 and 2.24) with high BLUP_SD values (1.67 and 1.45), suggesting unstable performance across seasons (Table 3.2).

**Table 3.2:** Genotype Selection Table Based on BLUP and BLUP_SD.

| Rank | Accession Name | BLUP<br>(Yield<br>Potential) | BLUP_SD<br>(Stability) | Interpretation |
| --- | --- | --- | --- | --- |
| 1 | IITA-TMS-IBA180146 | 4.97 | 0.42 | High yield and stable, ideal for broad adaptation due to low BLUP_SD |
| 2 | IITA-TMS-IBA180081 | 3.50 | 0.78 | Moderate BLUP_SD, Good yield, moderately stable |
| 3 | IITA-TMS-IBA980581<br>(WChk) | 3.22 | 1.12 | High yield but responsive, season/environment-sensitive |
| 4 | IITA-TMS-IBA180210 | 2.65 | 0.39 | Moderate yield, highly stable due to low BLUP_SD |
| 5 | IITA-TMS-IBA180073 | 2.24 | 1.45 | Yield varies widely; season/environment-sensitive, high BLUP_SD |
| 6 | IITA-TMS-IBA180244 | 2.14 | 0.91 | Reliable with decent yield |
| 7 | IITA-TMS-IBA180017 | 2.01 | 0.33 | Modest yield, stable due to low BLUP_SD |
| 8 | IITA-TMS-IBA180049 | 1.79 | 1.38 | Low yield and unstable-less desirable, high BLUP_SD |
| 9 | IITA-TMS-IBA180065 | 1.58 | 0.72 | Low-yielding, moderate BLUP_SD |
| 10 | IITA-TMS-IBA180182 | 1.25 | 1.67 | Poor yield, unstable and least promising with high BLUP_SD and low BLUP |

#### Crossover Interaction was observed at 9 MAP

Crossover Interaction was noticed at 9-Months After Planting. Genotype rankings at this stage vary significantly between seasons, confirming crossover interaction. Genotypes like IITA-TMS-IBA180146 and IITA-TMS-IBA180081 ranked highest in 2020/2021 but were mid-ranked in 2019/2020. Conversely, IITA-TMS-IBA980581 (WChk) and IITA-TMS-IBA180244 were top-ranked in 2019/2020 but dropped substantially in 2020/2021. This rank reversal indicates that genotype performance is season-dependent, underscoring the need for multi-environment testing and season-specific selection in cassava breeding (Table 3.3).

**Table 3.3:**
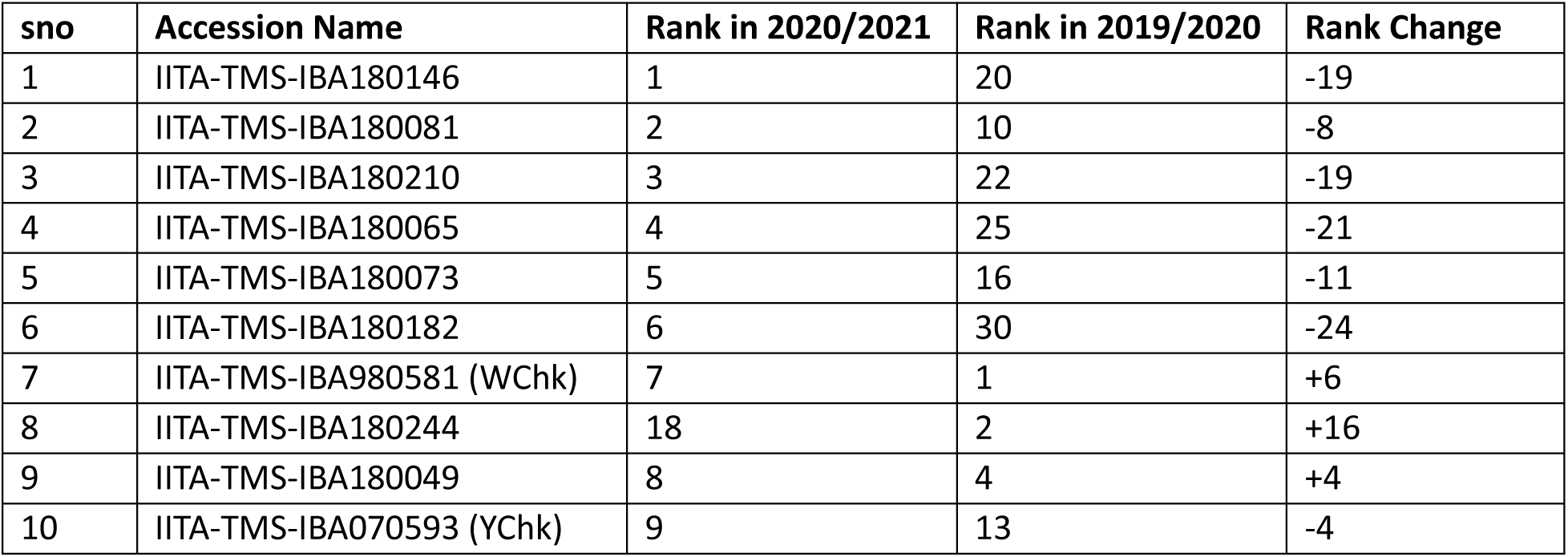
Genotypic Rank Changes Between Seasons at 9 Months After Planting.

#### Genotypic Performance and Stability Across Seasons via Heatmap and Cluster Analysis

A heatmap was generated to visualize the yield performance of cassava genotypes across two cropping seasons (2019/2020 and 2020/2021). Each row in the heatmap corresponds to a genotype, while each column represents a season. The color gradient-from white to deep red-reflects the magnitude of yield, with red indicating higher performance and white denoting lower yield.

This visual representation allows for rapid identification of genotypes with consistent or season-specific performance. Genotypes exhibiting deep red coloration across both seasons are considered high performers with stable yield, suggesting broad adaptability. In contrast, genotypes that shift from red in one season to white in another demonstrate crossover interaction, where performance is highly dependent on seasonal conditions. This observation aligns with earlier findings at MAP 9, where genotype rankings varied significantly between seasons (Figure 3.1).

**Figure3.1:**
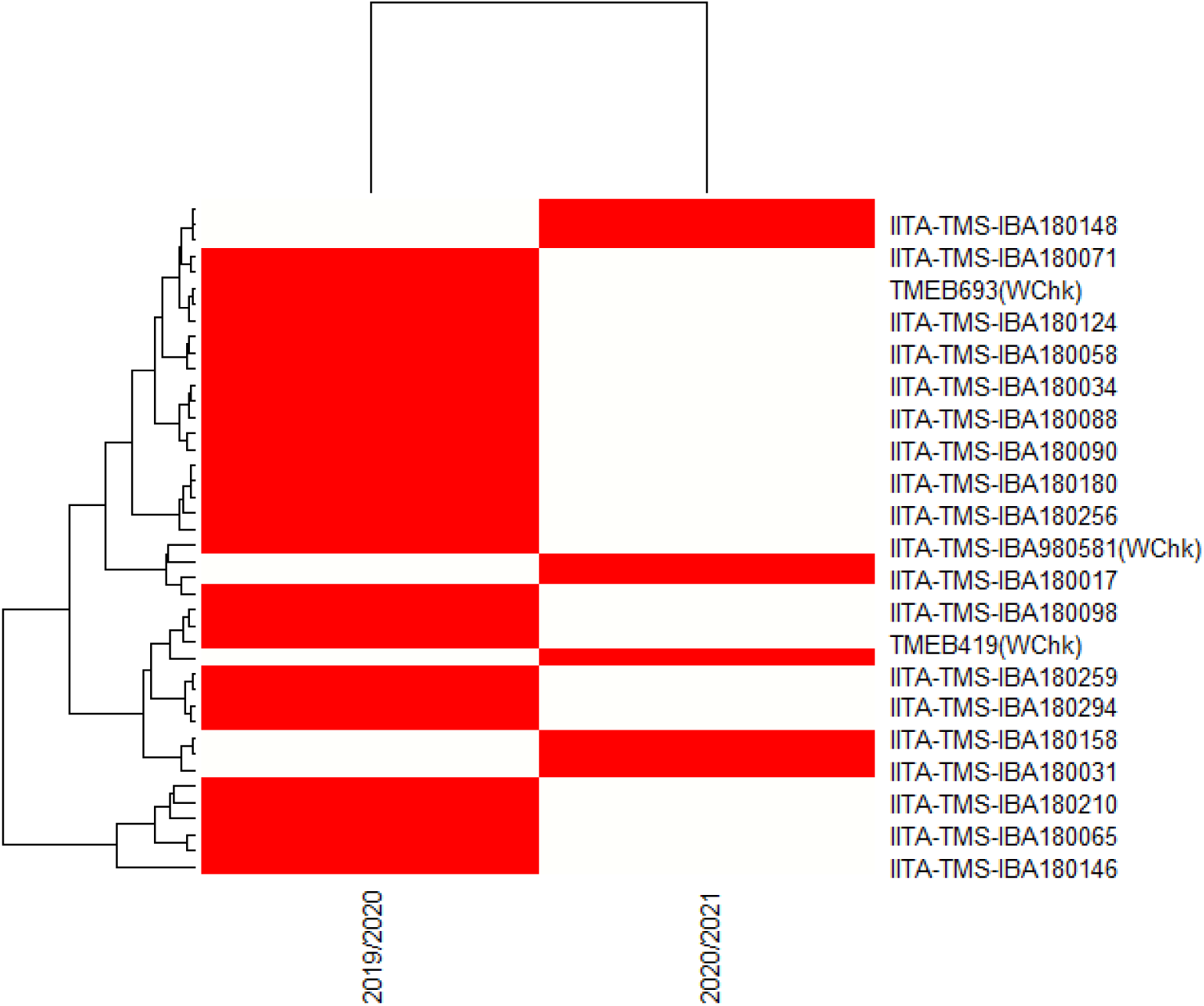
Hierarchical Clustering and Heatmap of Cassava Genotype Yield Performance Across Seasons. The heatmap displays BLUP-estimated yield values for cassava genotypes across the 2019/2020 and 2020/2021 seasons. Red indicates high yield performance, while white indicates low yield. The dendrogram groups genotypes based on similarity in seasonal performance, revealing crossover interactions and environment-specific responses.

#### Genotypic Performance Across Seasons from Heatmap Analysis

The heatmap provides a visual summary of genotype yield performance across two cropping seasons (2019/2020 and 2020/2021). Each row represents a distinct genotype, while each column corresponds to a specific season. The color gradient from yellow to deep red indicates the magnitude of yield, with deeper red tones reflecting higher performance and lighter shades indicating lower yield.

This visualization enables rapid identification of genotypes with consistent or season-specific performance. Genotypes that display intense red coloration across both seasons are considered stable high performers, suggesting strong adaptability and reliable productivity. In contrast, genotypes that show red in one season and yellow or pale tones in another exhibit crossover interaction, where performance varies depending on environmental conditions. Such genotypes may be responsive to specific seasonal factors but lack broad stability (Figure 3.2).

**Figure3. 2:**
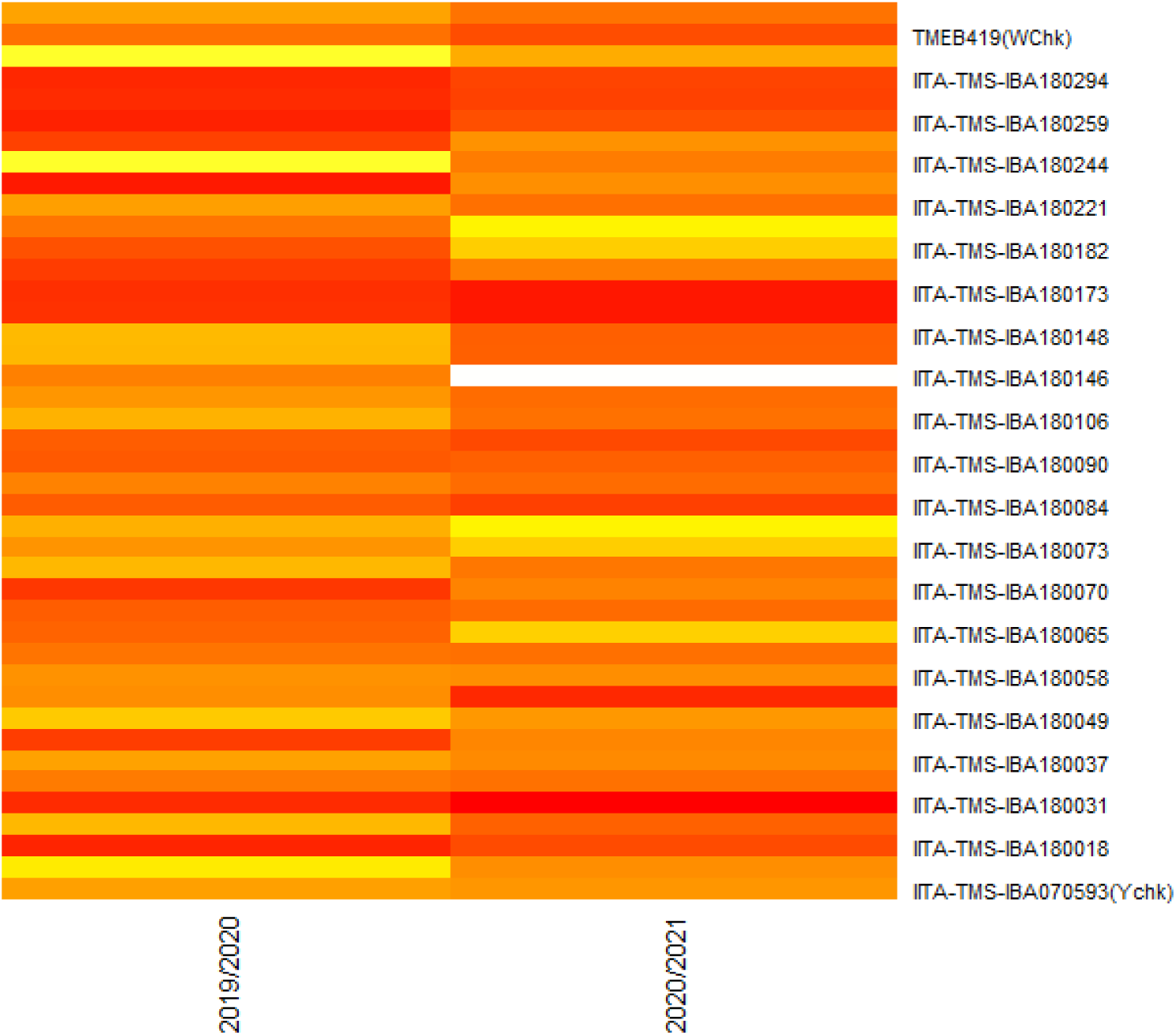
Heatmap of yield performance of genotypes across two cropping seasons.

### Model 4

**Table 4.1:** Variance Components from Model 4.

| Random Effect | Variance | Std. Dev | % of Total Variance | Interpretation |
| --- | --- | --- | --- | --- |
| MAP | 63.65 | 7.97 | 19.84% | Major environmental influence |
| Seasons | 18.86 | 4.34 | 5.88% | Moderate impact across years |
| Accession_name | 10.71 | 3.27 | 3.34% | Genotypic contribution relatively small |
| REP | 0.0002 | 0.013 | 0% | Differences between replicates within environments had minimal impact on yield. |
| Residual | 227.54 | 15.08 | 70.94% | Unexplained variance dominates |
| Heritability (H²) | 0.033 |  |  | A modest genetic contribution to FYLD. |

#### Residual and MAP Effects Account for Majority of Yield Variance

The variance components derived from the random-effects model used to partition sources of variation in fresh root yield is as presented in table 4.1. The largest proportion of variance (70.98%) was attributed to residual effects, indicating that unexplained environmental factors or experimental noise dominated yield variability. Among the structured components, MAP (Months After Planting) contributed the most (19.82%), confirming its strong environmental influence. About 6% of the yield differences were due to changes between seasons, showing that the variation from year to year was noticeable but not substantial. Genotypic variance, represented by differences among accessions, was relatively small (3.32%) and corresponds directly to the estimated broad-sense heritability (H²), indicating limited genetic differentiation and minimal genetic control over FYLD under the tested conditions. The fixed-effect intercept was not statistically significant (p = 0.116), consistent with the model’s emphasis on random effects. Overall, these results underscore the predominance of environmental factors over genetic contributions in shaping cassava yield performance.

#### Yield Variation Driven Primarily by Random Environmental Factors

The summary of fixed effects from the final variance partitioning model is presented in Table 4.2. The intercept estimate (12.32) represents the baseline fresh root yield across all environments and genotypes. However, this estimate was not statistically significant (p = 0.116), likely due to high variability across environments. No other fixed effects were included in this model, as the primary goal was to quantify the contribution of random effects particularly rainfall (MAP), seasonal variation, and genotypic differences to overall yield variation. These results confirm that the majority of modeled variation was captured by the random effects, with MAP and residual variance being the dominant contributors.

**Table 4.2:** Fixed Effects Summary.

| Effect | Estimate | Std. Error | df | t-value | p-value | Significance |
| --- | --- | --- | --- | --- | --- | --- |
| Intercept | 12.32 | 5.95 | 3.77 | 2.07 | 0.111 | ns |

ns-not significant

#### Proportion of Variance Attributed to Random Effects in Fresh Root Yield

The bar chart as illustrated in Figure 4.1 shows the relative contributions of different random effects-MAP (Months After Planting), Seasons, Accession_name, REP and Residual-to the total phenotypic variance in fresh root yield. The Residual component dominates, accounting for approximately 71% of the variance, indicating substantial unexplained variability or environmental noise. MAP contributes nearly 20%, reinforcing its role as a major environmental driver of yield. Seasonal effects explain about 6%, while genotypic variation (Accession_name) accounts for only 3%, suggesting limited genetic differentiation among accessions under the tested conditions.

This visualization complements Table 4.1 by clearly depicting the disproportionate influence of environmental factors over genetic contributions in cassava yield performance. Despite accounting for key environmental and genetic factors, over 70% of the total phenotypic variance in fresh root yield remained unexplained, as indicated by the dominant residual component in Model 4. This suggests that additional unmeasured variables such as soil nutrient profiles, microclimatic fluctuations, pest and disease pressure, and subtle management practices may have influenced yield outcomes. The limited genotypic contribution (3.32%) further underscores the complexity of yield expression under field conditions. Future studies should consider integrating spatial data, soil diagnostics, and biotic stress monitoring to better capture the multifactorial nature of cassava productivity.

**Figure 4.1:**
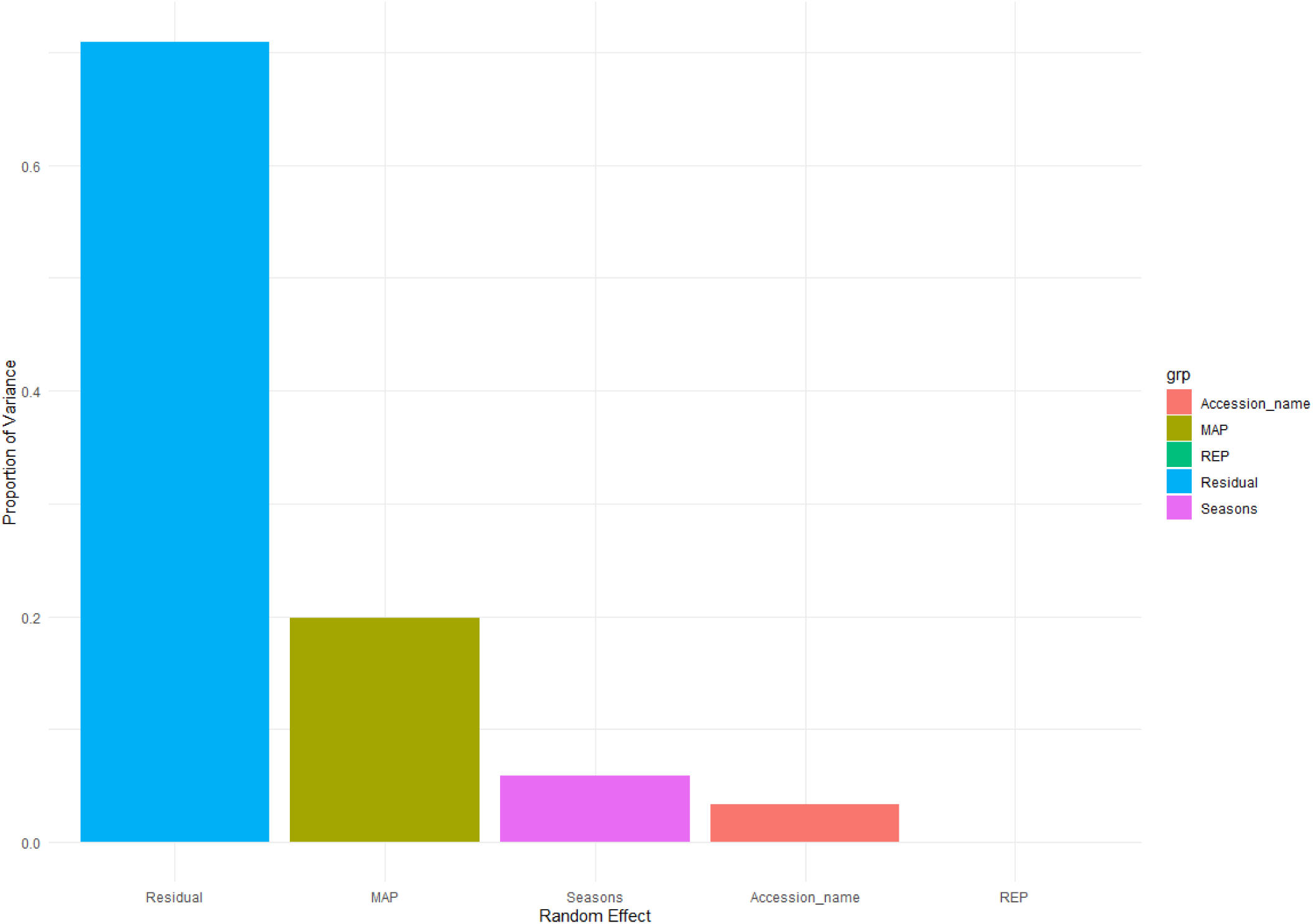
Proportion of Environmental Variance in Fresh Root Yield.

#### Climatic Variability Reinforces Seasonal Effects on Cassava Yield

As shown in Figure 4.2, the bar chart compares total rainfall (in millimeters) between the two growing seasons: 2019/2020 and 2020/2021. The 2019/2020 season recorded substantially higher rainfall, exceeding 100 mm, while the 2020/2021 season experienced markedly lower precipitation, below 20 mm. This stark contrast in seasonal rainfall highlights the environmental variability across trial years and may help explain differences in genotype performance and residual variance observed in the yield models (Figure 4.2). This figure supports the interpretation of seasonal effects in Model 4, reinforcing the role of climate variability in shaping cassava productivity.

**Figure 4.2:**
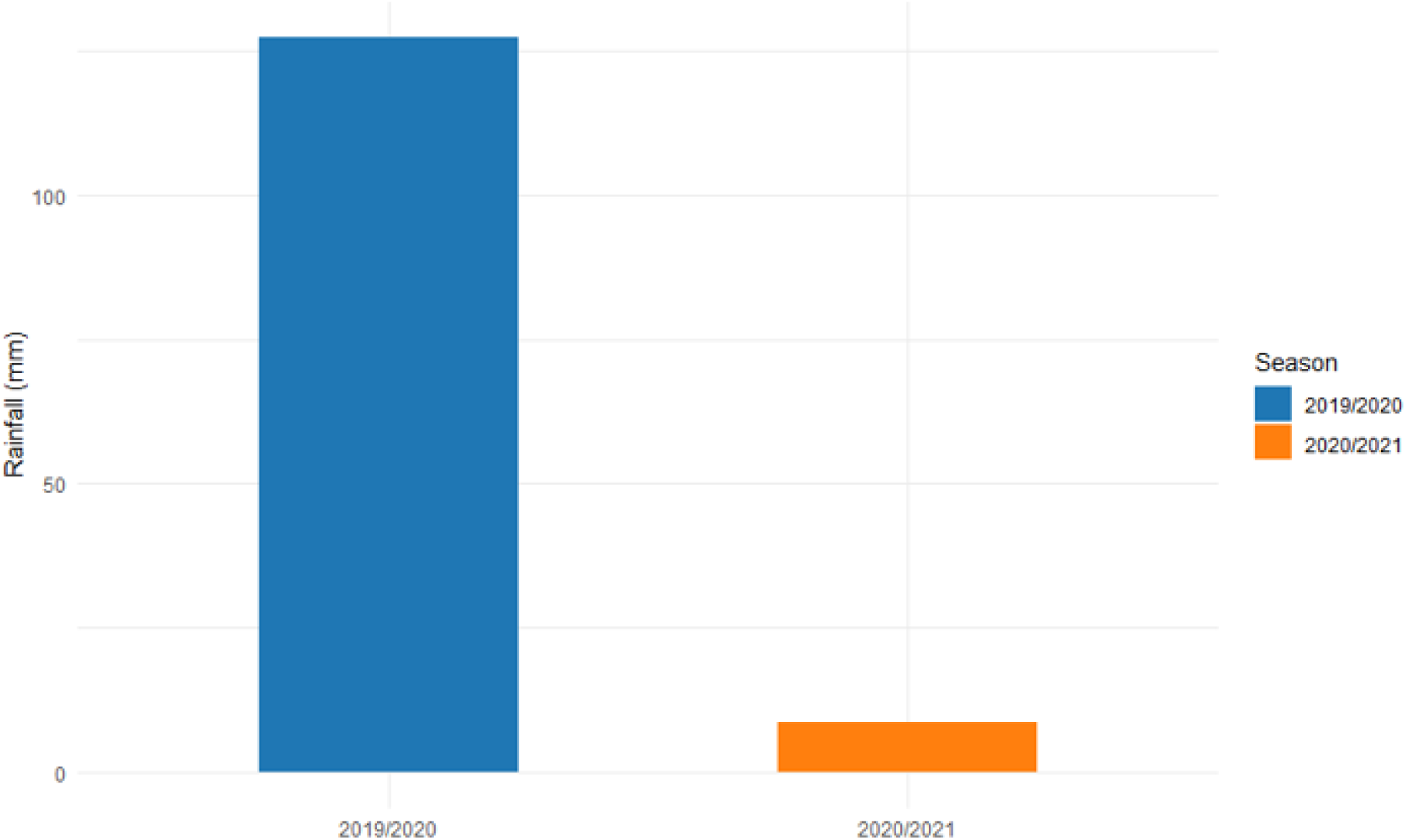
Seasonal Rainfall Comparison.

#### Moisture Availability at Harvest Stages Influences Genotypic Performance

This bar chart illustrates rainfall variation across three harvest periods defined by Months After Planting (MAP): 6MAP (December), 9MAP (March), and 12MAP (June). Rainfall was highest at 12MAP, exceeding 90 mm, followed by 9MAP with moderate rainfall around 45 mm, while 6MAP recorded minimal precipitation, just above 0 mm.

These seasonal rainfall patterns provide environmental context for interpreting yield variability. The low rainfall at 6MAP may have contributed to reduced root development or stress-induced performance in some genotypes, while the higher moisture availability at 12MAP likely supported optimal growth. This figure reinforces the importance of MAP as a key environmental driver, as highlighted in Models 1 and 4 (Figure 4.3).

**Figure 4.3:**
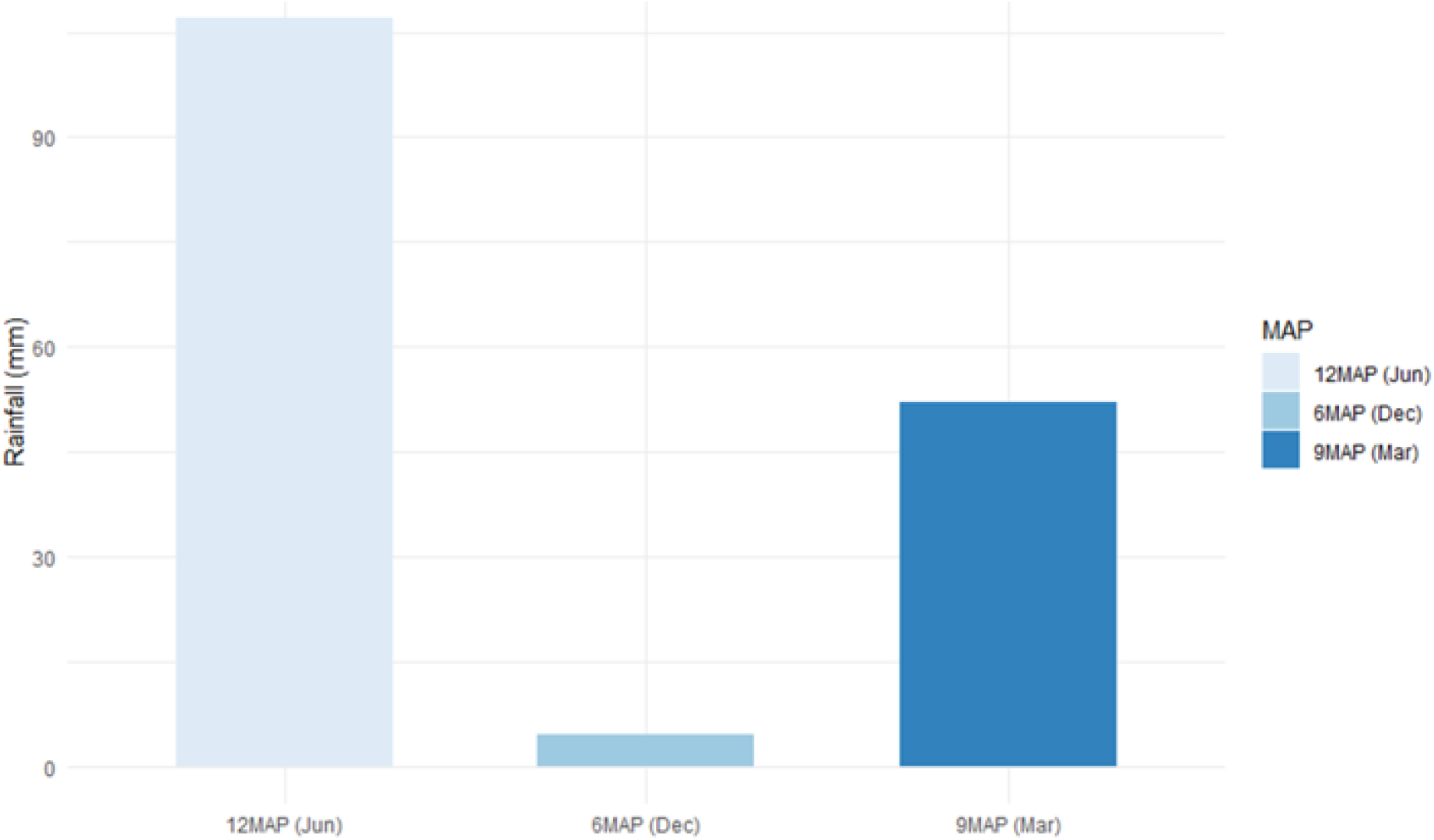
Rainfall Across MAP.

## Discussion

Given the polygenic nature of cassava yield shaped by numerous small-effect loci and complex genotype × environment interactions, no single model can fully capture the breadth of variation observed in field trials (Okogbenin et al., 2013). This study employed a combination of fixed-effect and mixed-effect models, each selected to illuminate different dimensions of yield variation. BLUPs and rank shift analyses further revealed genotype-specific responses, particularly under drought stress, which were not fully captured by fixed-effect models alone. Although residual variance remained high across models, this reflects the inherent complexity of cassava yield and the influence of unmeasured micro-environmental factors (Phanthanong et al., 2025; Sampaio Filho et al., 2024). Using different models and checking them carefully helped ensure that the results are both statistically reliable and meaningful, rather than depending on just one type of analysis (Burnham & Anderson, 2002).

## Model 1-Baseline Mixed Model (BLUP)

### Yield Variation Explained by MAP Alone

#### Dominance of Environmental Influence

The statistical significance of MAP (F = 92.54, p <0.001) underscores its overwhelming impact on cassava yield (Badewa et al., 2020). Neither Seasons nor REP contributed meaningfully to variability in Fresh Root Yield (FYLD), and the MAP × Seasons interaction was negligible. The results imply that cassava yield is more strongly shaped by site-specific environmental factors than by seasonal variation or replication design.

### Genotypic Performance via BLUPs

#### BLUPs revealed accessions with high genetic contributions to yield

IITA-TMS-IBA180146, IBA980581 (WChk), and IBA180081 consistently ranked among the top-performing genotypes, exhibiting high BLUPs for fresh root yield and stable performance across environments. These results suggest strong overall genetic merit and potential suitability for selection under prevailing agroecological conditions. In contrast, accessions such as IBA180031, IBA180158, and IBA180173 showed consistently negative BLUPs, indicating limited yield potential and poor adaptation to the test environments (Mbe et al., 2024).

The shrinkage property of BLUPs pulls extreme predictions toward the population mean, lending robustness to selection under variable environments. This approach is ideal for identifying candidates with consistently superior performance across trials (Melchinger et al., 2024). Genotypes exhibiting high BLUPs for fresh root yield can be prioritized for advancement to multi-location trials or inclusion in recommendation lists, given their strong overall genetic merit and environmental stability. Conversely, accessions with consistently low BLUPs may warrant reassessment or exclusion from further evaluation due to limited performance and poor adaptation

The estimated heritability of 4.4% signals a low genetic contribution to observed variation in cassava yield (Olasanmi et al., 2022). This low heritability estimate (H²) is not uncommon for complex traits such as fresh root yield, especially in early-stage trials or under conditions of high environmental variability. It aligns with the ANOVA results, where Months After Planting (MAP) emerged as the only highly significant factor, suggesting that genotype performance was driven more by developmental timing than by inherent genetic differentiation.

Under these conditions, selection precision is limited, as phenotypic performance shows a weak association with underlying genotypic variation. To improve reliability in identifying superior accessions, multi-environment trials may be necessary to detect genotypes whose performance is minimally influenced by environmental fluctuations

## Model 2-Fixed-Effect Genotypic Model

### MAP and Seasonal Effects on Yield

#### Genotype-Level Performance

By incorporating both MAP and season, this model captures how environmental fluctuations interact with developmental stages. Seasonal rainfall variability emerges as a key driver of yield shifts, especially in later MAP intervals.

Accessions IITA-TMS-IBA180146 once again emerged as a top performer, reinforcing its suitability for deployment. IITA-TMS-IBA180146 shows strong and consistent yield potential in both models. While its p-value is borderline, the magnitude of its fixed estimate and BLUP makes it a strong candidate for recommendation.

The three underperforming accessions not only have strongly negative fixed effects but are also statistically significant, and their BLUPs further confirm poor genotypic performance. These may warrant reconsideration in selection pipelines. These accessions IITA-TMS-IBA180031, IITA-TMS-IBA180158, and IITA-TMS-IBA180173 had statistically significant negative yield effects, suggesting removal from elite pools or further evaluation under different conditions (Sampaio et al.,2023).

#### Environmental Effects

MAP was highly significant at P<0.001 and continued to show strong yield influence, while Seasons reached statistical significance and was highly significant (P<0.001) unlike in Model 1. This suggests accessions may respond differently depending on seasonal conditions when modeled as fixed effects (Sampaio et al., 2024).

#### Comparison with BLUPs

BLUPs and fixed estimates aligned directionally, confirming the robustness of genotype performance patterns. However, fixed effects were more sensitive to extremes, and unlike BLUPs, they allow formal hypothesis testing. Model 2 would be important if the objective was to directly evaluate particular accessions, especially in varietal trials or recommendation pipelines (Barbiero et al., 2024).

## Model 3-G×E Interaction Model

### Effects of Genotype, Environment, and Their Interactions on Yield Variation

#### Environmental Dominance Persists

The third model in this study examined the Genotype × Environment (G×E) interaction to assess differential genotypic responses across seasons and locations. Although the interaction terms were not statistically significant, the strong main effects of MAP and Seasons confirm that environmental factors play a dominant role in shaping yield outcomes.

G×E interactions in this study, were modeled as Genotype × Environment, where Environment was defined by the stratified combination of months after planting (MAP) and cropping seasons. This structure reflects changes over time and across different seasons, making it possible to observe how each genotype performs under different growth phases and weather conditions (Sampaio et al., 2024).

The absence of significant G×E effects suggests that genotype rankings were relatively stable across environments. However, visual tools such as rank shift tables and BLUP_SD (Standard Deviation of Best Linear Unbiased Prediction) analysis revealed crossover patterns that may still hold biological importance for breeding decisions (Sampaio et al., 2023).

### BLUP-Based Ranking of Cassava Genotypes with Associated Stability Indicators

The integration of BLUP and BLUP_SD offers a nuanced understanding of genotype performance beyond raw yield values. Genotypes with high BLUP and low BLUP_SD such as *IITA-TMS-IBA180146* are prime candidates for broad adaptation due to their consistent high performance across environments (Kartika et al., 2013). These genotypes are particularly valuable in breeding programs targeting yield stability under variable agroecological conditions.

Genotypes with high BLUP but elevated BLUP_SD-such as *IITA-TMS-IBA980581(WChk-White Check)* may be responsive to specific environmental cues, performing exceptionally well under favorable conditions but inconsistently elsewhere. While these genotypes hold promise for targeted deployment, their instability limits their utility in wide-scale cultivation.

Conversely, genotypes with low BLUP and low BLUP_SD such as *IITA-TMS-IBA180017* are reliable but agronomically less desirable due to limited yield potential. Genotypes with both low BLUP and high BLUP_SD such as *IITA-TMS-IBA180182* are the least promising, as they combine poor productivity with unpredictable performance. This dual-metric approach reinforces the importance of selecting genotypes that balance yield potential with environmental stability, ensuring both productivity and reliability in cassava improvement programs.

### Crossover GxE Interactions

#### Visual Detection of Crossover Genotype × Environment Patterns Despite Statistical Non-Significance

The genotype-by-environment interaction (G×E) was examined across two cropping seasons using BLUP estimates, heatmap visualization, and dendrogram clustering. Several genotypes exhibited notable shifts in performance between seasons, as reflected in the heatmap’s color transitions from deep red (high yield) to pale or white (low yield). These reversals were especially evident in genotypes such as *IITA-TMS-IBA180073* and *IITA-TMS-IBA180182*, which performed well in one season but poorly in another.

Notably, genotype rank shifts and crossover interactions were most evident at 9MAP, where intermediate rainfall conditions appeared to modulate genotypic responses, leading to performance reversals across environments (Mbe et al., 2024).

Such performance shifts are indicative of crossover interactions, where genotype rankings change depending on the season. These dynamics are not captured by global p-values from mixed models or ANOVA, which test for overall effects but may overlook rank reversals. Instead, season-specific BLUP comparisons and visual tools like heatmaps reveal these nuanced patterns.

The accompanying dendrogram illustrates hierarchical clustering of genotypes based on their yield profiles. Genotypes positioned on closely linked branches share similar response patterns across seasons, suggesting potential genetic or physiological similarities in environmental adaptation. These clusters offer valuable insight into genotype groupings that may respond similarly to agronomic conditions and can inform targeted breeding strategies.

Together, the heatmap and dendrogram provide a powerful means of evaluating both yield potential and stability. They highlight genotypes suited for broad adaptation as well as those better deployed in specific environments. Genotypes that appear red in one season and white in another demonstrate clear evidence of crossover interaction, underscoring their sensitivity to environmental factors.

This pattern reinforces the influence of G×E, suggesting that some genotypes are season-responsive rather than universally stable. While they may be valuable in targeted environments, their inconsistent performance across seasons limits their suitability for broad deployment. Breeding strategies should therefore consider both yield potential and stability, prioritizing genotypes that maintain high performance across diverse conditions.

The presence of large rank shifts among genotypes confirms the occurrence of crossover G×E interactions-a critical insight for breeding decisions. These interactions suggest that certain genotypes excel under specific seasonal conditions but underperform elsewhere, emphasizing the need for environment-matching in deployment.

These findings also highlight the limitations of relying solely on global significance tests. Breeding programs aiming for wide adaptation should prioritize genotypes with stable performance across seasons, while those targeting specific environments may benefit from selecting responsive genotypes with season-dependent strengths.

## Model 4-Variance Partitioning Model

### Residual Variance and the Limits of Current Predictors

#### Environmental Influence Outweighs Genotypic Effects in Yield Variation

Model 4 confirmed that cassava yield is overwhelmingly shaped by Months After Planting (MAP) and residual factors, not by genotype or season alone. The 20% contribution of MAP validates its consistent impact across all models. Meanwhile, genotypic variance remained low (3%), confirming earlier observations from Models 1 to 3. Breeders should prioritize matching genotypes to specific environmental conditions rather than relying solely on general genotype recommendations.

The bar plot highlights how MAP clearly dominates the structured variability (nearly 20%), while Genotype and Seasons contribute modestly. The large residual component (71%) reflects either high noise or potential G×E interactions not accounted for in this additive model while the replicate effect contributed negligibly to yield variation, as indicated by its near-zero variance (0.0002) and standard deviation (0.013), accounting for 0% of the total variance.

#### Unexplained Variance and Hidden Environmental Factors

Despite accounting for rainfall, MAP, and seasonal variation, a substantial proportion of residual variance remained across all models, particularly in Model 4, where residuals exceeded 70% of total phenotypic variation. This persistence signals the presence of unmodeled environmental variables or trial-specific nuances not captured by categorical or rainfall metrics alone. These potential contributors to unexplained variation may include Soil fertility (e.g., potassium, nitrogen levels), humidity, and temperature extremes, uneven field management practices (e.g., planting depth, weeding), pest and disease pressures, microclimate fluctuations or site-specific anomalies (Oliveira et al., 2012). These factors may be nested within broader MAP and Season categories, but remain statistically unresolved in their current form. Their influence warrants future inclusion as covariates or stratification layers to better resolve genotype performance under variable conditions.

#### Heritability and Selection Potential

The estimated broad-sense heritability (H² = 0.033) was modest, indicating low efficiency of direct phenotypic selection for fresh root yield under the tested conditions. This low heritability may reflect the inherent complexity of yield as a trait, limited replication, environmental uniformity, or substantial residual variation from unmeasured micro-environmental factors (e.g., soil nutrient levels, field stressors). In such contexts, marker-assisted selection or genomic prediction may offer complementary strategies to enhance selection accuracy for yield-related traits, especially when genetic control is weak

#### Impacts of Rainfall on Cassava Performance

To deepen the understanding of agroecological and seasonal influences on cassava yield, observed rainfall data were analyzed across months after planting (MAP) and trial seasons. The environmental contrast was striking: total rainfall dropped from 127.38 mm in the 2019/2020 season to just 8.45 mm in 2020/2021, placing the latter within an extreme drought context. Similarly, rainfall varied sharply across MAP, ranging from only 4.58 mm at 6MAP (dry season) to 107.05 mm at 12MAP (wet season).

These patterns align closely with the variance component analysis, which identified rainfall (MAP) as the most influential structured factor, accounting for nearly 20% of the total variance in fresh root yield. The pronounced environmental contrast between seasons and planting stages likely amplified the observed variability in yield, while the low heritability estimate (H² = 0.033) suggests that genetic differences among accessions were largely masked by these environmental extremes. This reinforces the conclusion that cassava yield performance in this study was predominantly shaped by rainfall availability and other unmeasured environmental stressors, rather than by genotypic potential alone.

#### This environmental partitioning adds critical explanatory power to the findings across all four models

In Models 1 and 2, MAP consistently emerged as a strong predictor of yield, now clearly linked to moisture availability. The seasonal significance observed in Model 2 can be biologically attributed to the stark rainfall disparity between years. In Model 3, genotypes such as IBA180146 and IBA180182 exhibited sharp rank shifts, which are interpretable as drought sensitivity under low-rainfall conditions. Meanwhile, the high residual variance in Model 4 likely reflects the impact of the severe drought in 2020/21, underscoring that broad categories like season and MAP alone are insufficient to fully capture the environmental complexity influencing yield.

## Limitations and Future Directions

While this study successfully dissected genotypic and environmental contributions to cassava yield using four robust mixed models, notable unexplained variation remained, particularly the high residual variance observed in Model 4 (>70%). This suggests that key environmental and agronomic variables influencing yield were not captured within the current MAP, Season, and rainfall parameters. The absence of these factors constrains model explanatory power and may obscure genotype-specific sensitivities or adaptations. Their inclusion as covariates or nested random structures in future trial designs is essential for improving precision in phenotypic prediction and trait dissection.

## Recommendations

The discovery of genotype × environment interactions particularly across Months After Planting (MAP)-highlights the need for targeted selection strategies. Rather than treating yield as a static trait, breeders should identify accessions that perform reliably at specific developmental stages. Some genotypes may exhibit early vigor while others show late-yield accumulation, making stage-specific evaluation essential for strategic deployment.

Given MAP’s strong influence on phenotypic variance, future field trials should be designed to evaluate cassava performance at key growth stages. Structured assessments at consistent intervals such as 6, 9, and 12 months after planting can help reveal how yield develops over time and how flexible each genotype is. Aligning evaluations with these developmental milestones will give breeders a clearer picture of each genotype’s potential and adaptability.

Expanding genotype evaluations across diverse agroecological zones and planting seasons would enhance the detection of stability and adaptive responses. Multi-location trials capture broader environmental variability and provide better resolution of genotype × environment effects. This approach is particularly valuable for identifying broadly adapted genotypes versus niche performers.

In light of the low genetic control over total yield, emphasis should be placed on more heritable component traits such as root size, harvest index, or canopy architecture. These traits may indirectly influence overall yield and offer more reliable pathways for genetic gain. Refining selection criteria to include physiologically meaningful traits can help stabilize performance and accelerate improvement.

The low heritability observed suggests that selection based solely on yield may be less effective, as the trait is predominantly influenced by non-genetic factors. To better isolate genetic effects and reduce residual noise, replicated trials across diverse environments are essential. Moreover, while MAP, Seasons, and Genotype contributed to yield variation, over 70% of the total phenotypic variance remained unexplained, as revealed by the dominant residual component in Model 4. This underscores the need to incorporate additional features into future models such as soil nutrient profiles, pest and disease pressure, planting density, and microclimatic data to more accurately capture the complexity of field conditions. Including these agronomic and environmental variables could significantly enhance model precision and deepen our understanding of the multifactorial drivers of cassava yield variability.

## Conclusion

This study applied a multi-model method to evaluate cassava genotypic performance and dissect the environmental and genetic drivers of fresh root yield. Models 1 and 2 consistently identified high-and low-performing genotypes, with IITA-TMS-IBA180146 emerging as a top candidate for selection. Model 3 confirmed the stability of genotypic rankings across seasons, indicating minimal crossover interactions. Model 4 revealed that environmental factors particularly rainfall timing (MAP) contributed more to yield variation than genetic differences, with residual variance accounting for over 70% of the total.

These findings underscore the complexity of cassava yield expression under field conditions and highlight the need for environment-specific breeding strategies. Genotypes such as IITA-TMS-IBA980581 and IITA-TMS-IBA180017 demonstrated strong responsiveness to high rainfall, suggesting their suitability for wetter agroecologies. Conversely, drought-sensitive accessions like IITA-TMS-IBA180146 and IITA-TMS-IBA180182 may require targeted improvement or exclusion from dry-season trials.

Future breeding efforts should integrate rainfall responsiveness, soil diagnostics, and biotic stress monitoring into selection pipelines. By refining environmental characterization and expanding trait evaluation, cassava improvement programs can enhance yield stability and resilience under increasingly variable climatic conditions.

